# Promoter capture drives the emergence of proto-genes in *Escherichia coli*

**DOI:** 10.1101/2023.11.15.567300

**Authors:** Md. Hassan uz-Zaman, Simon D’Alton, Jeffrey E. Barrick, Howard Ochman

## Abstract

The phenomenon of *de novo* gene birth—the emergence of genes from non-genic sequences—has received considerable attention due to the widespread occurrence of genes that are unique to particular species or genomes. Most instances of *de novo* gene birth have been recognized through comparative analyses of genome sequences in eukaryotes, despite the abundance of novel, lineage-specific genes in bacteria and the relative ease with which bacteria can be studied in an experimental context. Here, we explore the genetic record of the *Escherichia coli* Long-Term Evolution Experiment (LTEE) for changes indicative of “proto-genic” phases of new gene birth in which non-genic sequences evolve stable transcription and/or translation. Over the time-span of the LTEE, non-genic regions are frequently transcribed, translated and differentially expressed, thereby serving as raw material for new gene emergence. Most proto-genes result either from insertion element activity or chromosomal translocations that fused pre-existing regulatory sequences to regions that were not expressed in the LTEE ancestor. Additionally, we identified instances of proto-gene emergence in which a previously unexpressed sequence was transcribed after formation of an upstream promoter. Tracing the origin of the causative mutations, we discovered that most occurred early in the history of the LTEE, often within the first 20,000 generations, and became fixed soon after emergence. Our findings show that proto-genes emerge frequently within evolving populations, persist stably, and can serve as potential substrates for new gene formation.

## Introduction

New genes are thought to originate mostly through a process of duplication and divergence, in which copies of already existing genes are repurposed to serve new functions [1–3]. This process, however, requires the presence of pre-existing genes and does not address how genes that serve as substrates for duplication and divergence originally arose. The *de novo* origin of genes from non-genic sequences, *i.e.*, regions other than existing RNA or protein-coding genes, began receiving consideration in the 1990s [4]. The potential for new genes to emerge in this way was reinforced by both the functional characterization of putative *de novo* genes [5,6] and the widespread identification of lineage-specific genes in virtually every species [7–12]. Furthermore, transcriptome surveys established that non-genic regions of the genome are continually subject to stochastic transcription and translation [13–16], such that new genes could arise if a non-genic sequence manifests more operative expression and a beneficial function.

*De novo* gene formation, which has been studied extensively in eukaryotes [17] and viruses [18,19], appears also to contribute to bacterial evolution. A substantial fraction of the gene repertoires of bacterial species are lineage-specific [20,21] and such genes often show no clear homologs, indicating that they may not have originated by duplication and divergence [20,22] or horizontal gene transfer [23,24]. Moreover, some lineage-specific genes have been traced to non-coding sequences, suggesting the possibility of *de novo* emergence [25]. Although bacterial genomes contain little intergenic DNA, both strands of their genomes are transcribed pervasively [26,27], and a number of bacterial genes have been found to be derived from the opposite strand or within shifted reading frames of existing genes [28–30].

*De novo* emergence of a new gene can be conceptualized as the co-occurrence of the following: (*i*) transcription of the sequence, and in the case of protein-coding genes, (*ii*) acquisition of a novel open reading frame (ORF) and (*iii*) translation of the ORF, and (*iv*) appearance of a beneficial function [31,32] (Figure 1A). Following the terminology offered in [32], non-coding sequences arriving at intermediate stages on the way to becoming new genes are considered “proto-genes”, represented by the transition of ancestrally silent sequences to a state of transcription or translation (Figure 1B).

**Fig 1.**
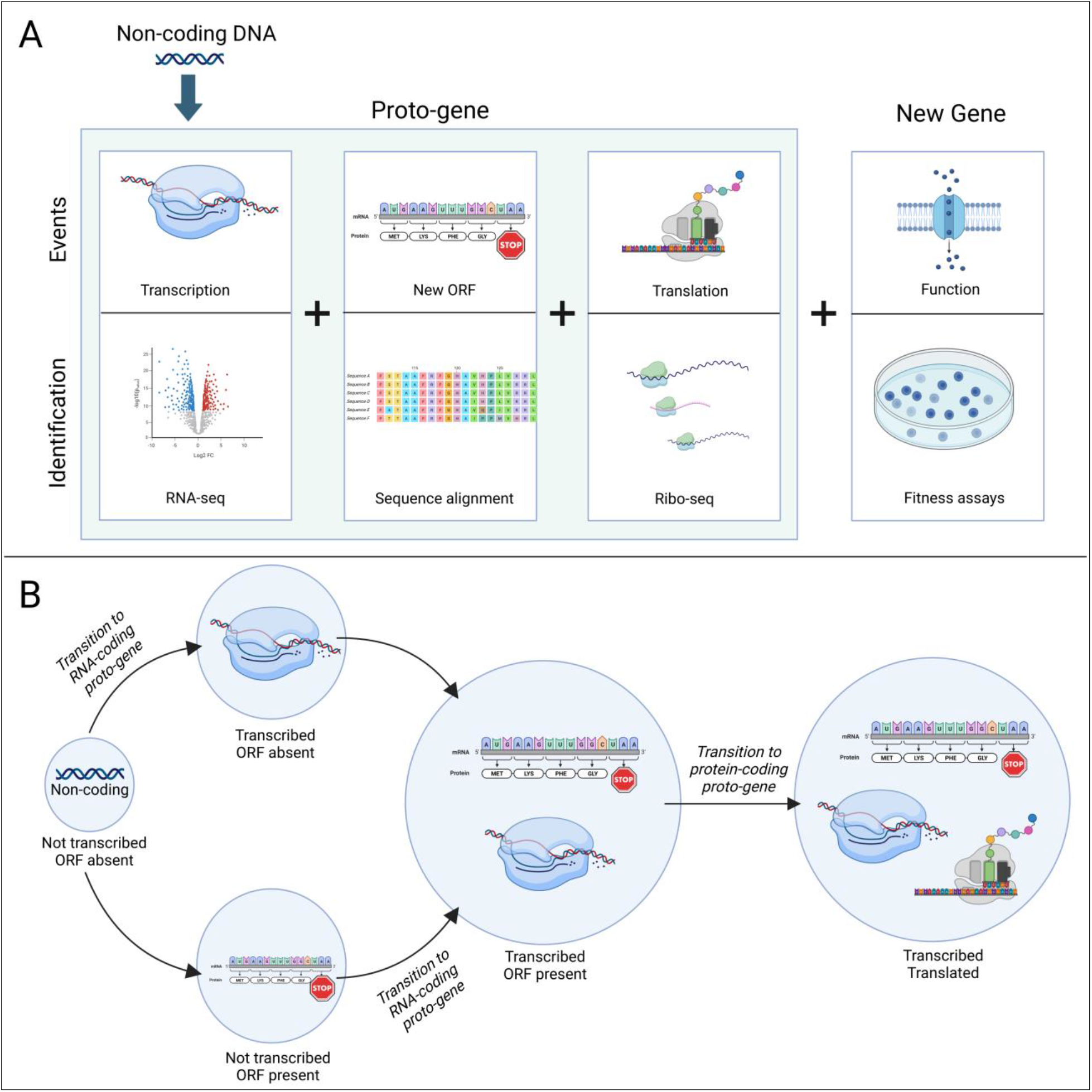
Stages of proto-gene emergence. (A) Events required for the birth of proto-genes (shaded area) and new genes, and examples of experimental methods that can identify these events. Modified and expanded from [31]. (B) Ways in which a non-coding region can transition into a proto-gene. Created with Biorender.

To date, most *de novo* genes have been recognized through retrospective and comparative analyses [33,34]; however, unicellular eukaryotes and bacteria offer the opportunity to experimentally investigate gene birth on account of their short generation times and potential for rapid evolution. In this context, the genome sequencing [35], transcriptomics and ribosome profiling [36] data generated for the *E. coli* Long-Term Evolution Experiment (LTEE) [35,37] allow direct detection of newly emerged proto-genes by following changes in expression and associated mutations in evolving lineages. We demonstrate that within the timescale and environment of the LTEE, proto-genes emerge most frequently via promoter capture, persist indefinitely once they arise, and reach fixation within the population.

## Results

### Non-genic transcription and translation are frequent in the LTEE

To assess the overall extent of transcription and translation occurring in non-genic regions in the LTEE, we surveyed genome-wide expression levels in 400-bp sliding windows along both strands of the genome. Of 18,801 such windows in the LTEE ancestor genome, 10,006 overlapped annotated genes on the same strand, 8,642 on the opposite strand (antisense), and 153 were intergenic (Supplementary File 1). Transcription and translation in non-genic regions, which include both antisense and intergenic windows, were detected in all lines and timepoints surveyed (Figure 2AB, Figure S2, Supplementary File 1).

We first examined RNA-seq data of clones isolated from 11 LTEE lines at 50,000 generations. In this dataset, 95.0% of windows overlapping annotated protein- or RNA-coding genes and 64.0% of the non-genic windows were transcribed in at least one clone at a relaxed threshold of 1 transcript per million reads (TPM). When raising the threshold to >5 TPM, 73.9% annotated and only 10.3% of non-genic windows met this more stringent cutoff. After eliminating regions located upstream or downstream of annotated genes (*i.e.*, within 100 bp on the same strand) to account for possible transcription readthrough, this fraction fell to 7.6% for non-genic windows (Supplementary File 1).

Focusing next on Ribo-seq data from the same clones, whereas a similar number of annotated windows were translated at the >1 TPM level as were transcribed (93% *vs*. 95%), the fraction of translated non-genic windows fell to 37.7%. At the more stringent cutoff, only 4.4% of non-genic windows contained reads >5 TPM (2.7% after eliminating regions with possible readthrough) compared to 71.9% of annotated windows. Among the more highly transcribed (>5 TPM) non-genic windows without readthrough (termed “isolated windows” in Figure 2), 25.1% met the >5 TPM Ribo-seq cutoff, whereas a vast majority (94.0%) of annotated windows transcribed at this level showed evidence of translation.

**Fig 2.**
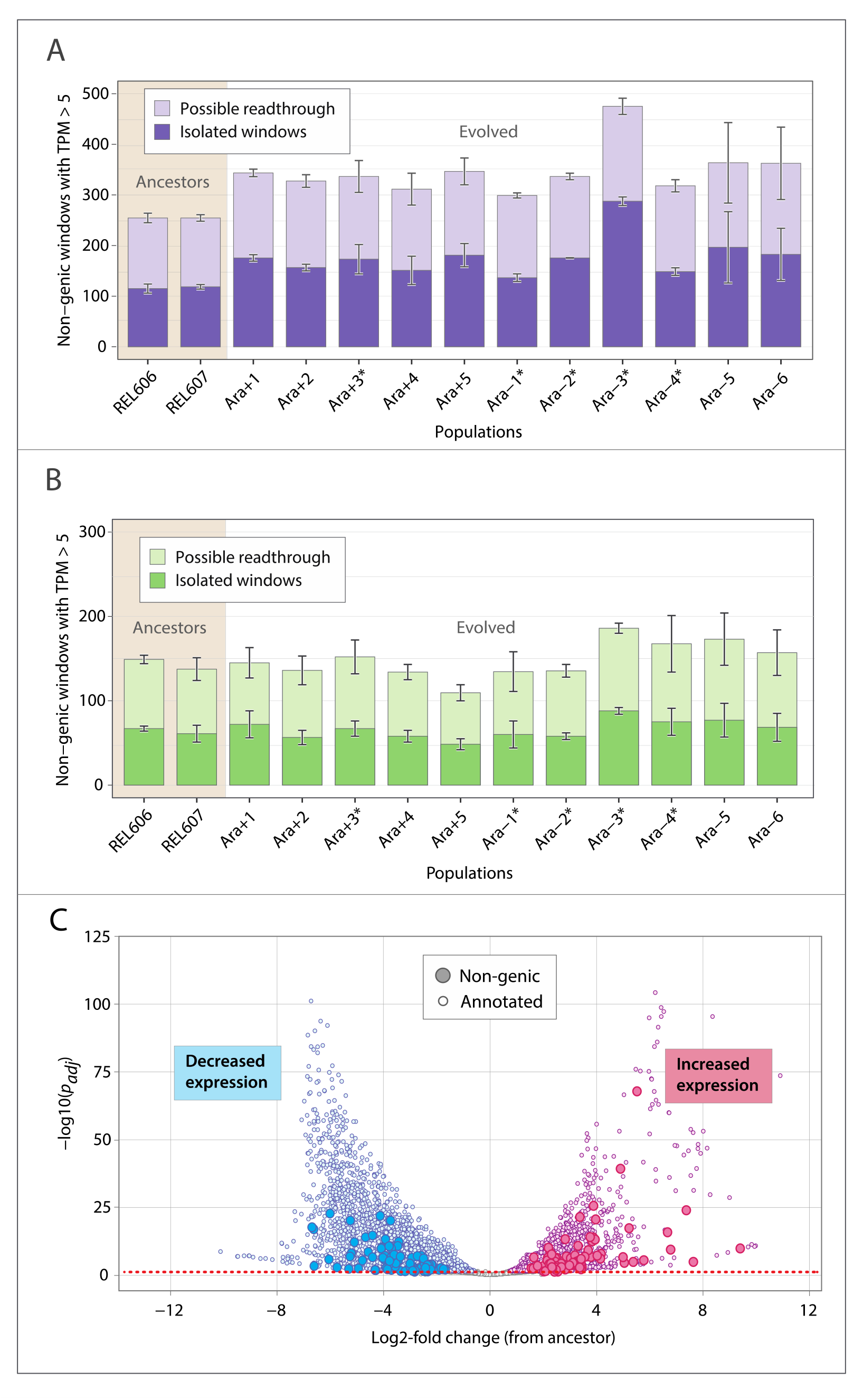
Expression of non-genic regions at 50,000 generations. Numbers of non-genic windows in ancestral (tan-shaded background) and evolved populations with normalized (A) RNA-seq and (B) Ribo-seq read counts expressed in average number of transcripts per million (TPM) across two biological replicates. Windows situated within 100 bp of an annotated gene are labelled as “Possible readthrough”, and those more than 100 bp from an annotated gene are labelled as “Isolated windows”. (C) Volcano plot of all windows whose expression changed between ancestral and evolved populations. Dotted red line denotes *padj* = 0.05.

Ancestral and evolved clones averaged 254.5 and 347.3 non-genic windows transcribed at >5 TPM, respectively. Regardless of transcription threshold, annotation category or mutation rate of populations, evolved clones had significantly higher numbers of transcribed windows than their ancestors (independent two-sample t-test, *p* < 0.05), although this trend was not observed for translation (Figure 2, Table S1). Furthermore, whereas a total of 139 non-genic windows experienced a statistically significant change in transcription at 50,000 generations compared to ancestors (*padj* < 0.05, Wald test *p*-value adjusted by Benjamini-Hochberg method) (Figure 2C), no non-genic window was found to be differentially translated after accounting for transcription change and readthrough. Taken together, the extent of transcription, translation and differential expression in non-genic regions of evolved genomes suggests the presence of substantial raw material for new gene formation in the LTEE.

### New mutations coincide with novel transcription

The non-genic transcription gains observed in the LTEE could be the consequence of biological noise or result from mutations leading to promoter acquisition. The latter is more likely to persist across generations and potentially be co-opted as new genes [38], so we focused on transcription increases coinciding with the appearance of new mutations to explore proto-gene emergence.

We first identified all cases in which a non-genic region immediately downstream of a new mutation experienced an increase in transcription compared to the ancestor, and there were 63 statistically significant cases (*padj* < 0.05, see Methods) in the 50,000- generation dataset. To specifically identify cases of novel transcription, we visually inspected expression coverage at each region to eliminate those that were transcribed at any level in the ancestor (Supplementary File 2). This procedure yielded 19 unambiguous proto-gene candidates with levels of increase spanning over two orders of magnitude (Figure 3). Since these regions can represent unannotated genes that were initially silent under the experimental conditions of the LTEE, we surveyed datasets that measured gene expression in the LTEE ancestor across a wide range of environmental conditions and growth phases [39,40]. After eliminating the candidates that exhibited transcription in any of the conditions (“non-LTEE transcription” in Figure 4), a final set of nine cases of novel proto-genes was identified at the 50,000-generation timepoint (Table 1, Table S2, Figure S3).

**Fig 3.**
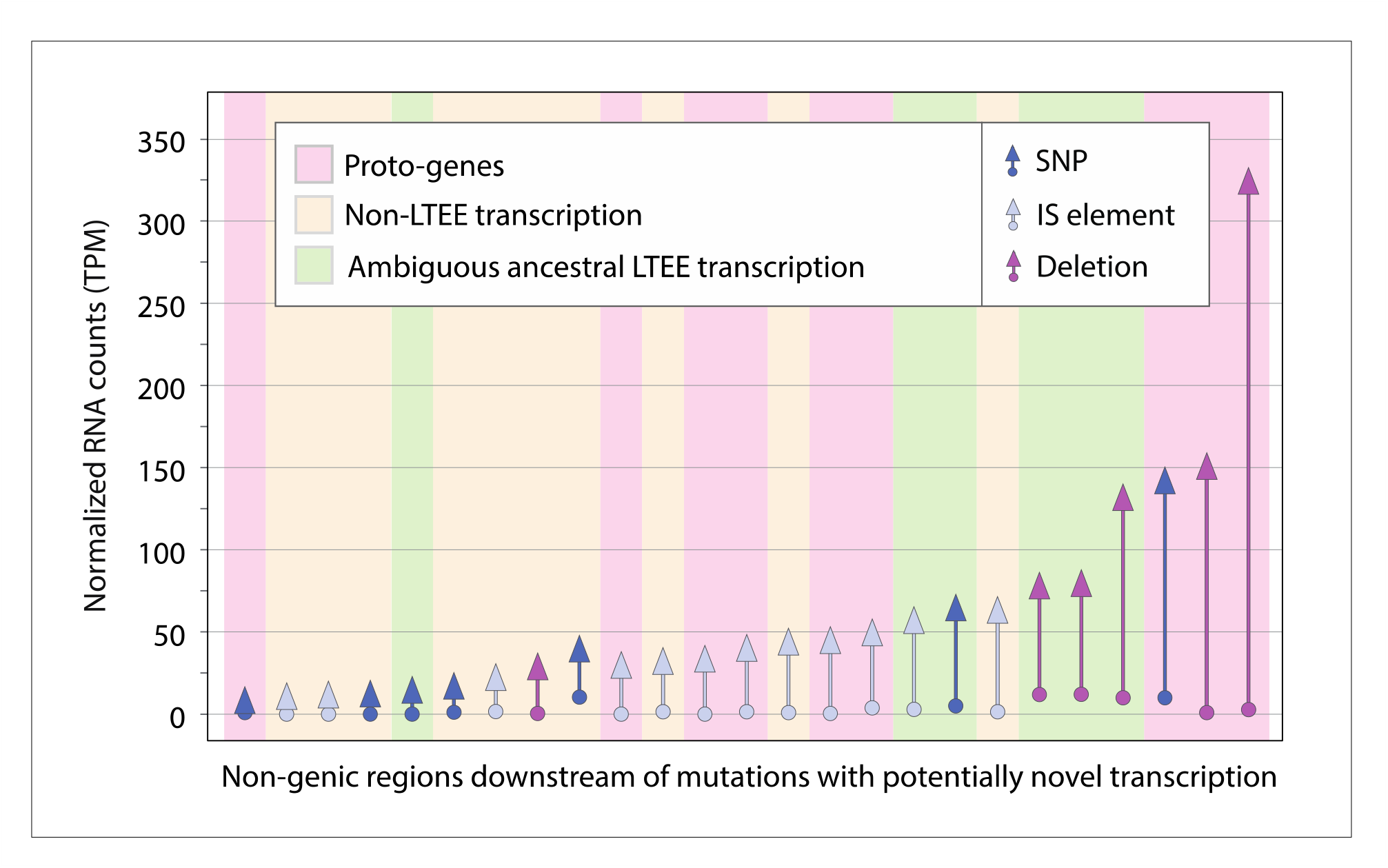
Mutations leading to novel transcription in LTEE populations. Arrows along *x*-axis represent regions displaying significant increases in expression, colored according to mutation type. Arrow lengths show differences in TPM between the ancestral and evolved states, with regions sorted according to the magnitude of change. Background shading represents different classes of proto-gene candidates.

**Fig 4.**
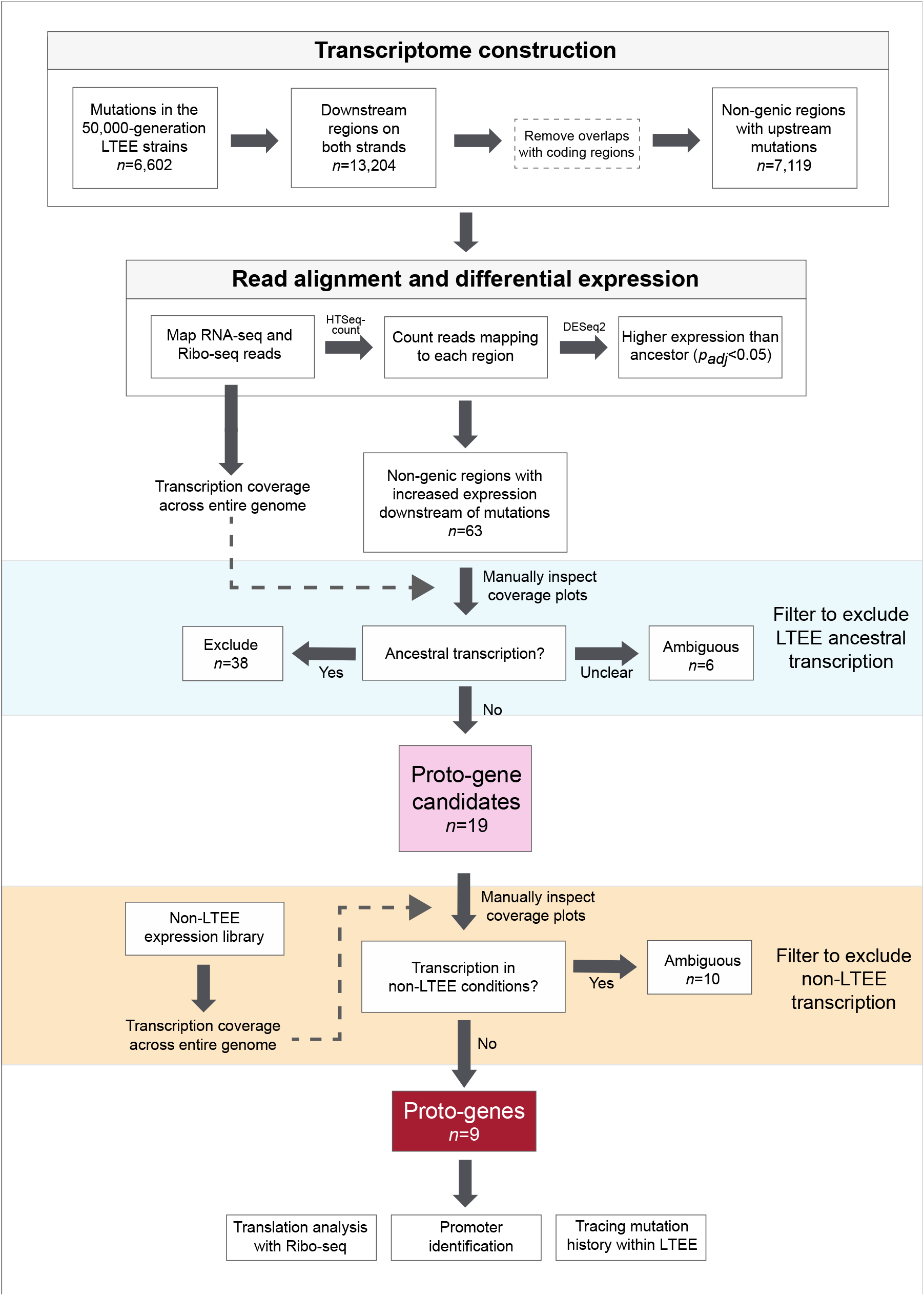
Workflow for detecting proto-genes.

**Table 1.**
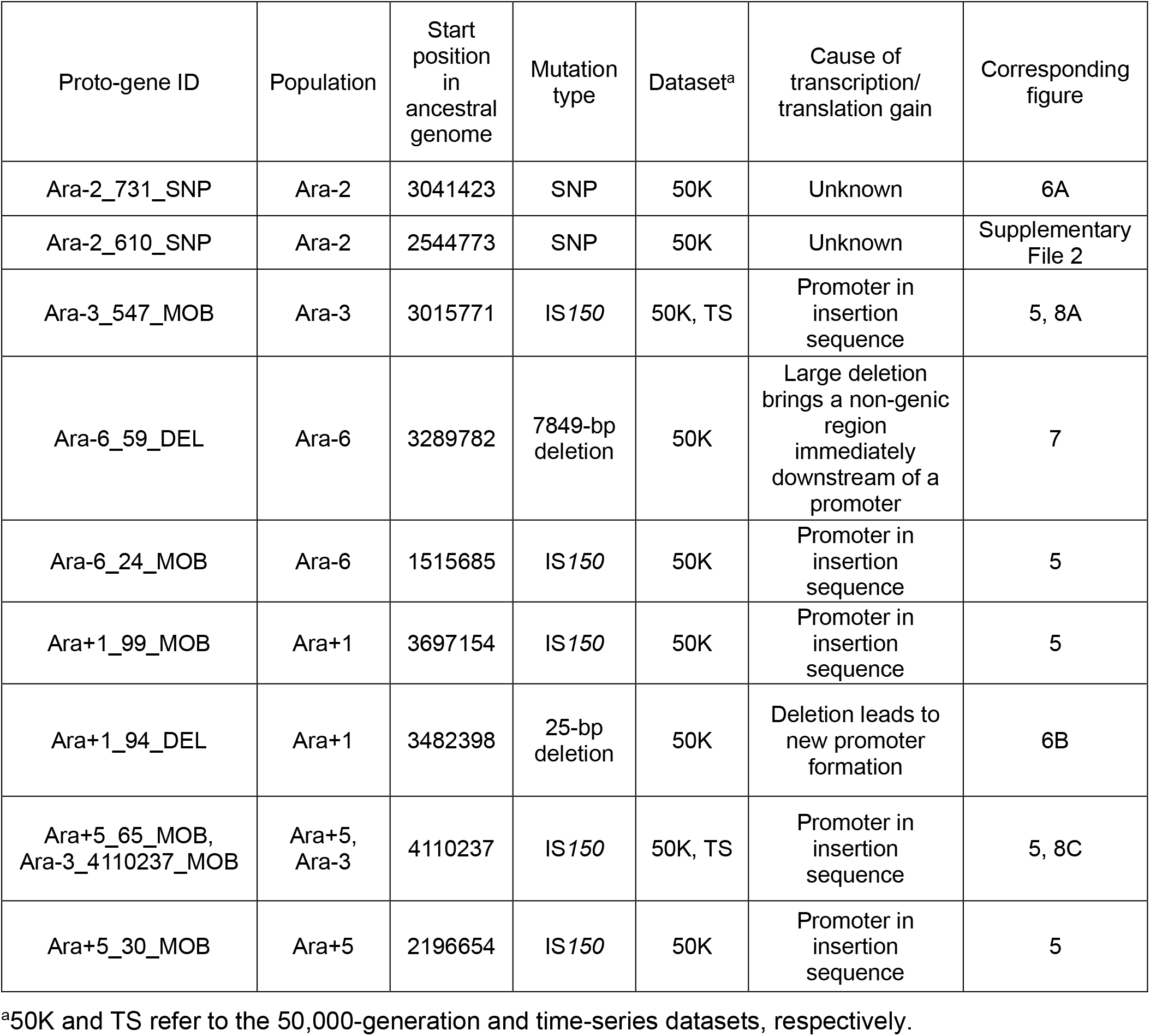
List of proto-genes identified in this study.

| Proto-gene ID | Population | Start position in ancestral genome | Mutation type | Dataset<sup>a</sup> | Cause of transcription/translation gain | Corresponding figure |
| --- | --- | --- | --- | --- | --- | --- |
| Ara-2_731_SNP | Ara-2 | 3041423 | SNP | 50K | Unknown | 6A |
| Ara-2_610_SNP | Ara-2 | 2544773 | SNP | 50K | Unknown | Supplementary File 2 |
| Ara-3_547_MOB | Ara-3 | 3015771 | IS150 | 50K, TS | Promoter in insertion sequence | 5, 8A |
| Ara-6_59_DEL | Ara-6 | 3289782 | 7849-bp deletion | 50K | Large deletion brings a non-genic region immediately downstream of a promoter | 7 |
| Ara-6_24_MOB | Ara-6 | 1515685 | IS150 | 50K | Promoter in insertion sequence | 5 |
| Ara+1_99_MOB | Ara+1 | 3697154 | IS150 | 50K | Promoter in insertion sequence | 5 |
| Ara+1_94_DEL | Ara+1 | 3482398 | 25-bp deletion | 50K | Deletion leads to new promoter formation | 6B |
| Ara+5_65_MOB, Ara-3_4110237_MOB | Ara+5, Ara-3 | 4110237 | IS150 | 50K, TS | Promoter in insertion sequence | 5, 8C |
| Ara+5_30_MOB | Ara+5 | 2196654 | IS150 | 50K | Promoter in insertion sequence | 5 |

<sup>a</sup>50K and TS refer to the 50,000-generation and time-series datasets, respectively.

To confirm that these 9 proto-genes are unique to the LTEE, we searched their sequences against a catalogue of both annotated and non-annotated transcripts assembled for *E. coli* K-12 MG1655 [41] consisting of 9,581 transcripts extracted from 3,376 RNA-seq experiments. All but one (Ara-3_547_MOB) of the nine proto-genes were present in the K-12 MG1655 genome, but none matched any annotated or unannotated transcript. The similarity between the LTEE ancestor and K-12 MG1655 (ANI = 99% [42]) indicates that expression of these sequences in *E. coli* prior to the initiation of the LTEE was unlikely, rendering them as *bona fide* instances of proto-gene emergence (Table 1).

### Insertion elements are frequent contributors to novel transcription

For five of the nine proto-genes, the alteration in their transcription activity was attributable to the activity of insertion sequence IS*150* (Figure 5), which carries an outward-facing promoter that can trigger downstream transcription. To investigate the generality of this effect, we surveyed all non-genic regions throughout the transcriptome that acquired an upstream IS*150* element by generation 50,000. Of 55 such regions, 12 displayed statistically significant increases in transcription, and in all but one case, the regions were silent in the LTEE ancestor. While these 11 passed the initial filter for proto-gene detection (Figure 4), six were excluded from the final list owing to their ambiguous transcription in the non-LTEE samples. The remaining 43 regions were either transcribed ancestrally or did not show a statistically significant increase. In general, transcription gains mediated by the IS*150*-promoter are modest compared to those caused by other types of mutations (Figure 3).

**Fig 5.**
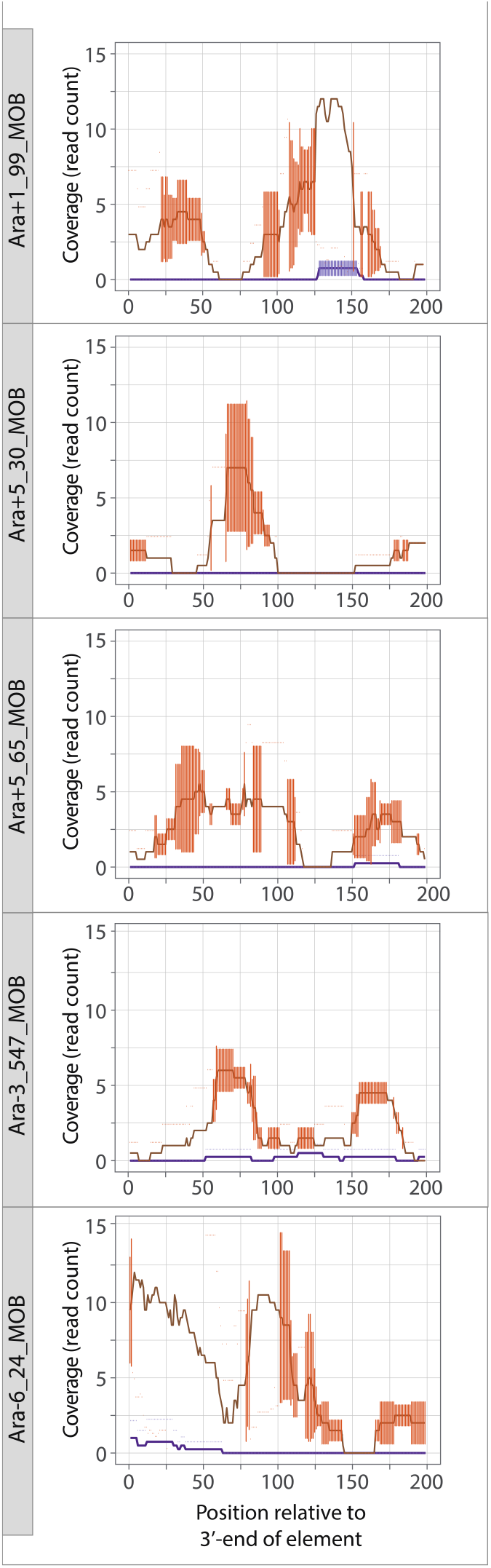
Novel transcription downstream of IS*150* insertions at generation 50,000. The five proto-genes formed by this mechanism are shown. Purple and orange lines denote ancestral and evolved transcription, respectively. Error bars represent standard deviation around the average value between replicates.

### Contribution of point mutations and small deletions to proto-gene emergence

Two of the proto-genes, both in the Ara-2 line, were putatively caused by upstream SNPs (Figure 6A), although newly formed promoters in the mutated regions were not recognized by promoter prediction tools [43,44]. In another case, in the Ara+1 line, the proto-gene was formed by a 25-bp deletion that brought two pre-existing sequences that resemble -10 and -35 promoter motifs into proximity, leading to the formation of a new promoter (Figure 6B). Although SNPs and small indels comprise the vast majority (95.7%) of the mutation-adjacent regions surveyed, these classes of mutations only rarely led to statistically significant transcriptional gains (Table S3).

**Fig 6.**
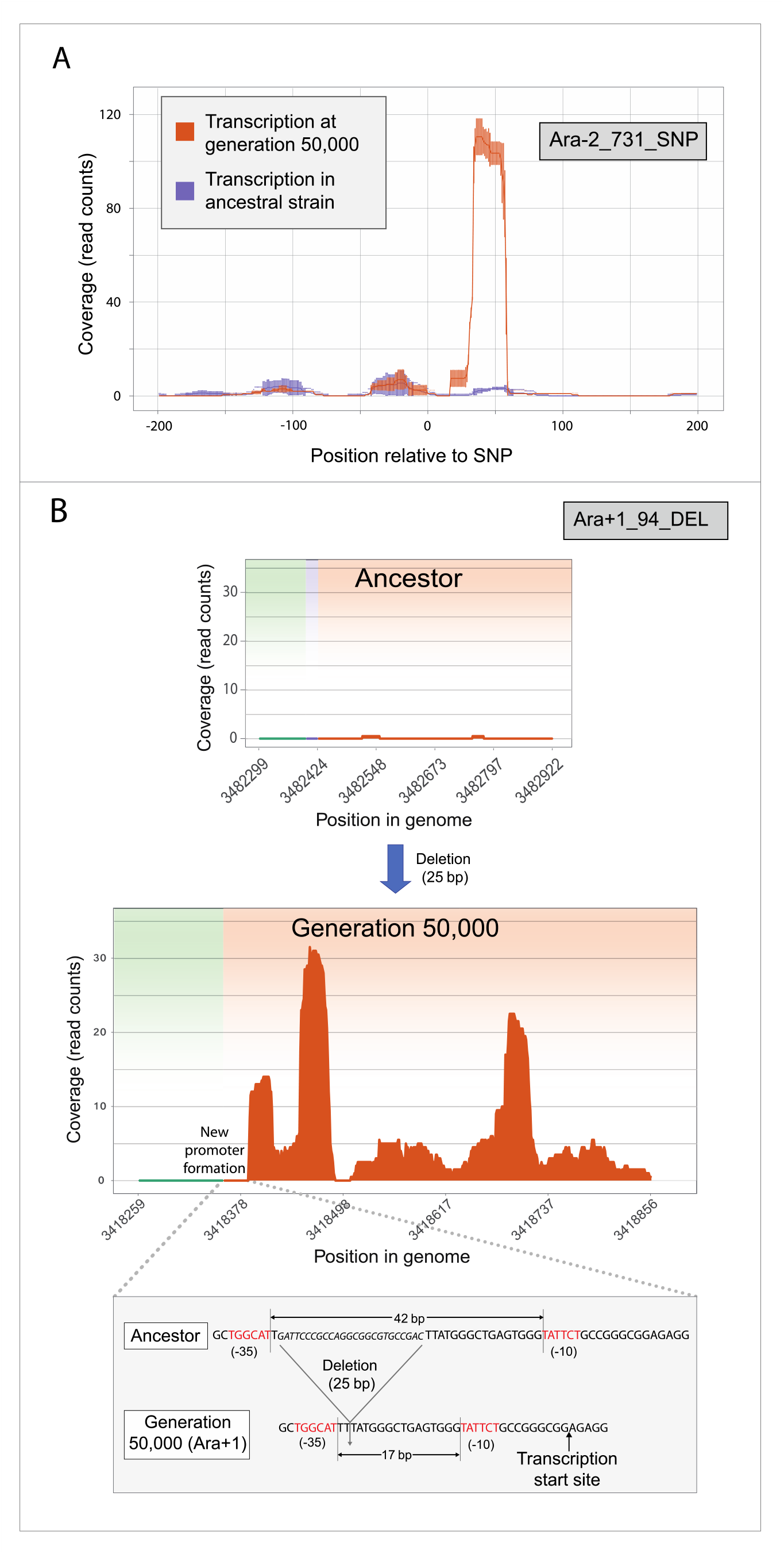
SNP- and deletion-associated expression of ancestrally non-transcribed regions. (A) SNP in Ara-2 lineage leading to downstream transcription, although no newly formed promoters were detected in this region. Error bars represent standard deviation around the average value between replicates. (B) A deletion of 25 bp in Ara+1 lineage led to *de novo* formation of a promoter and downstream expression. Green and orange shading represent regions immediately upstream and downstream of the mutation, respectively.

### Emergence of a peptide-encoding proto-gene via a large deletion event

With respect to the expression levels of the proto-genes, the largest increase was elicited by a 7849-bp deletion that overlapped six genes (*gltB, gltD, yhcG, ECB_03080, yhcH, nanK*) in the Ara-6 line (Figure 7A). This deletion shifted a transcriptionally silent region antisense to *nanK* and *nanE* to a position adjacent to and downstream of the promoter region of the glutamate synthase operon, resulting in new transcription. In the ribosome profiling data from this clone, there are pronounced buildups of reads in the corresponding region (Figure 7B), indicative of translated ORFs. The longest ORF (117 aa) is homologous to hypothetical proteins in *E. coli* and *Shigella sonnei*, indicating that it has gene-like features as recognized by standard prokaryotic annotation pipelines.

**Fig 7.**
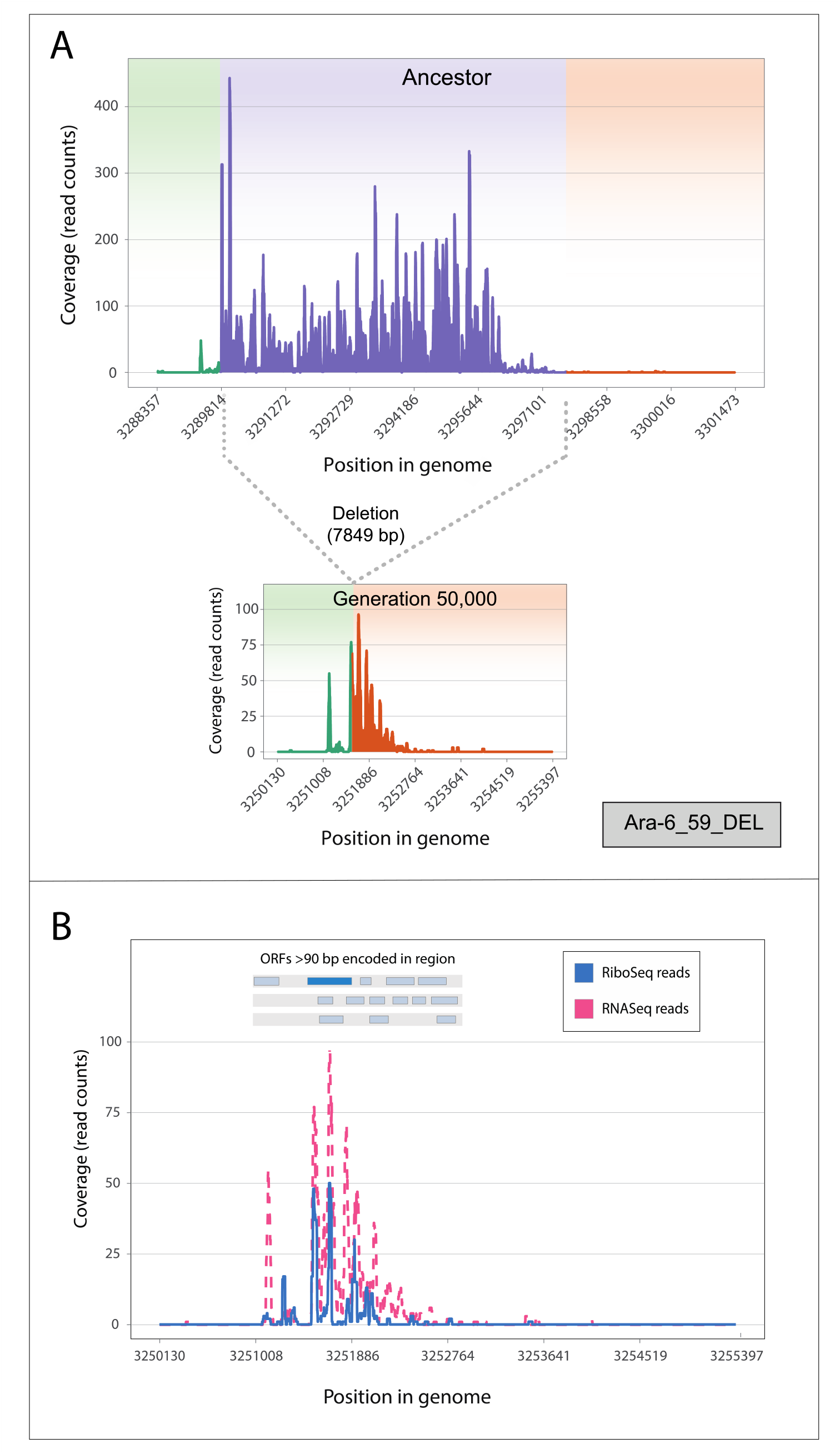
Large deletion-induced transcription and translation of ancestrally non-transcribed regions. (A) 7849-bp deletion spanning multiple genes (violet) placed a non-transcribed region in proximity of strong promoter (green), leading to expression downstream (orange). (B) Translation of same region in evolved population. Top insert shows positions of open reading frames >90 bp within region encoded on transcribed strand (dark blue segment denoting the longest ORF).

### Upregulation is stable across thousands of generations

If the proto-genes formed by new mutations at generation 50,000 were of recent origin and transitory, it would diminish the likelihood that these newly expressed regions would be co-opted as new genes. Therefore, to investigate the extent to which transcriptional changes forming proto-genes persist in the LTEE lines, we performed a complementary analysis to detect proto-genes using a time-series RNA-seq dataset of the Ara-3 line (Figure 4, S1). We found nine candidates with significantly increased expression at any evolved timepoint, of which five showed no transcription in the original LTEE ancestor (Supplementary File 3, Table S2). Two of these five were also recognized in the 50,000-generation data for the same line (above), with one retained in the final list of proto-genes (Table 1). In addition, we detected a new proto-gene in the time-series dataset that was not recognized in the 50,000-generation dataset because its expression change did not reach statistical significance. Interestingly, this proto-gene arose independently in the Ara+5 line via the same mutation. Overall, these results demonstrate that newly emerged proto-genes can persist indefinitely in the LTEE (Figure 8).

**Fig 8.**
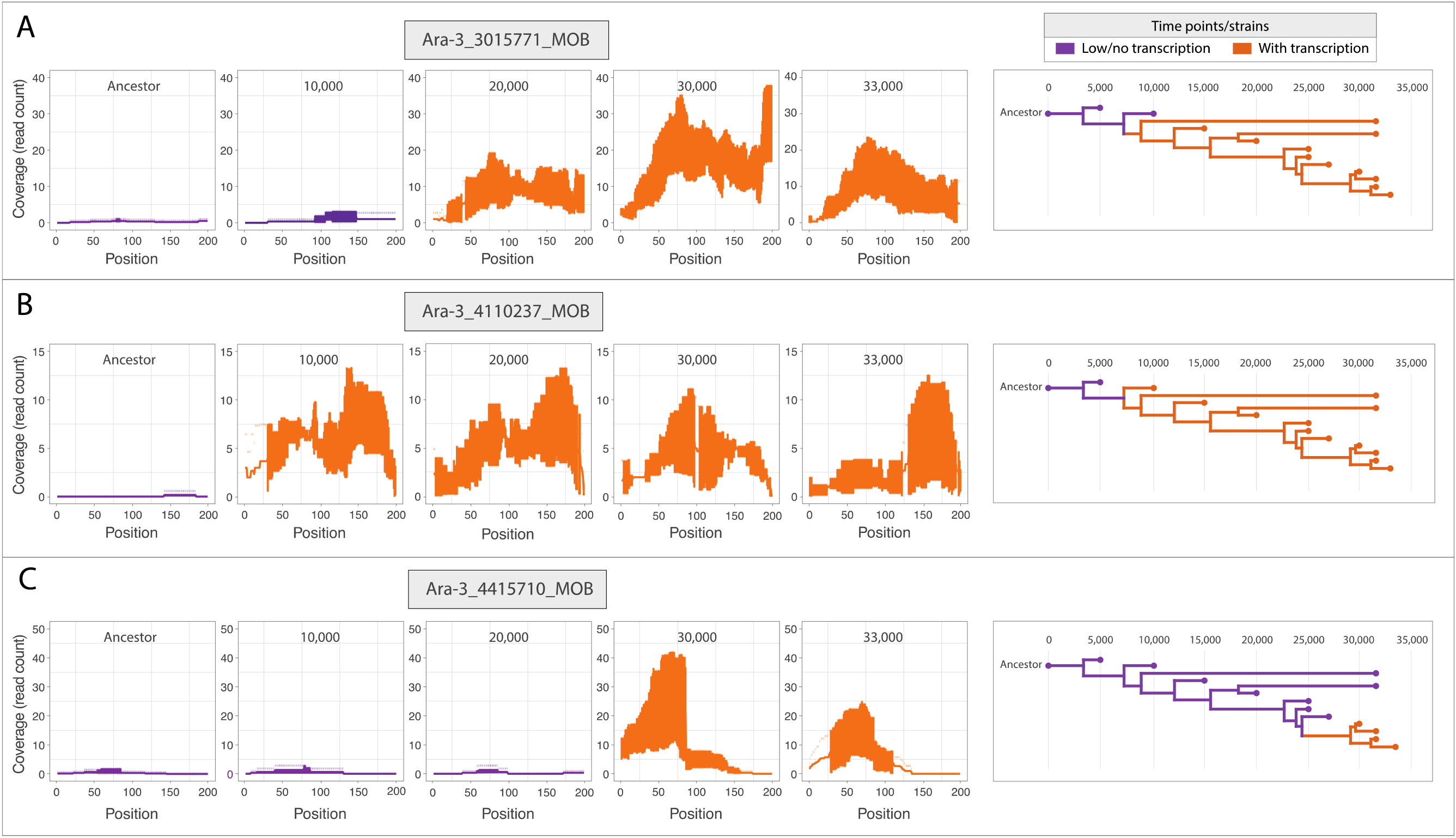
Persistent gains in transcription in Ara-3 population. Left: Levels of transcription in three proto-genic regions (A, B, C) at five timepoints in the LTEE. Error bars represent standard deviation around the average value between replicates. Right: Genealogies of time-series clones showing trasnscription status of corresponding regions at each time point.

### Proto-genes can arise rapidly and get fixed in evolving populations

To further investigate the persistence of proto-genes, we traced the origins of their associated mutations. To this end, we searched for the 10 proto-gene-associated mutations (nine from the 50,000-generation dataset, and one from the time-series) in genomic data generated for the first 60,000-generations of the LTEE [35]. Leveraging this population-sequencing dataset, we inferred the frequency of each mutation across timepoints, and in all but one case, the mutation was abundant in the population at both earlier and later timepoints (Figure 9). The three proto-genes with the largest increases in transcription (Figure 6, Figure 7) arose before the 20,000-generation timepoint and became fixed soon thereafter. The Ara+5_65_MOB mutation, an IS*150* insertion in the Ara+5 line, was not observed at the 50,000-generation timepoint, and only occurred at low frequencies at two subsequent timepoints. This mutation occurred independently in the Ara-3 line, suggesting the existence of an insertional hotspot in this region.

**Fig 9.**
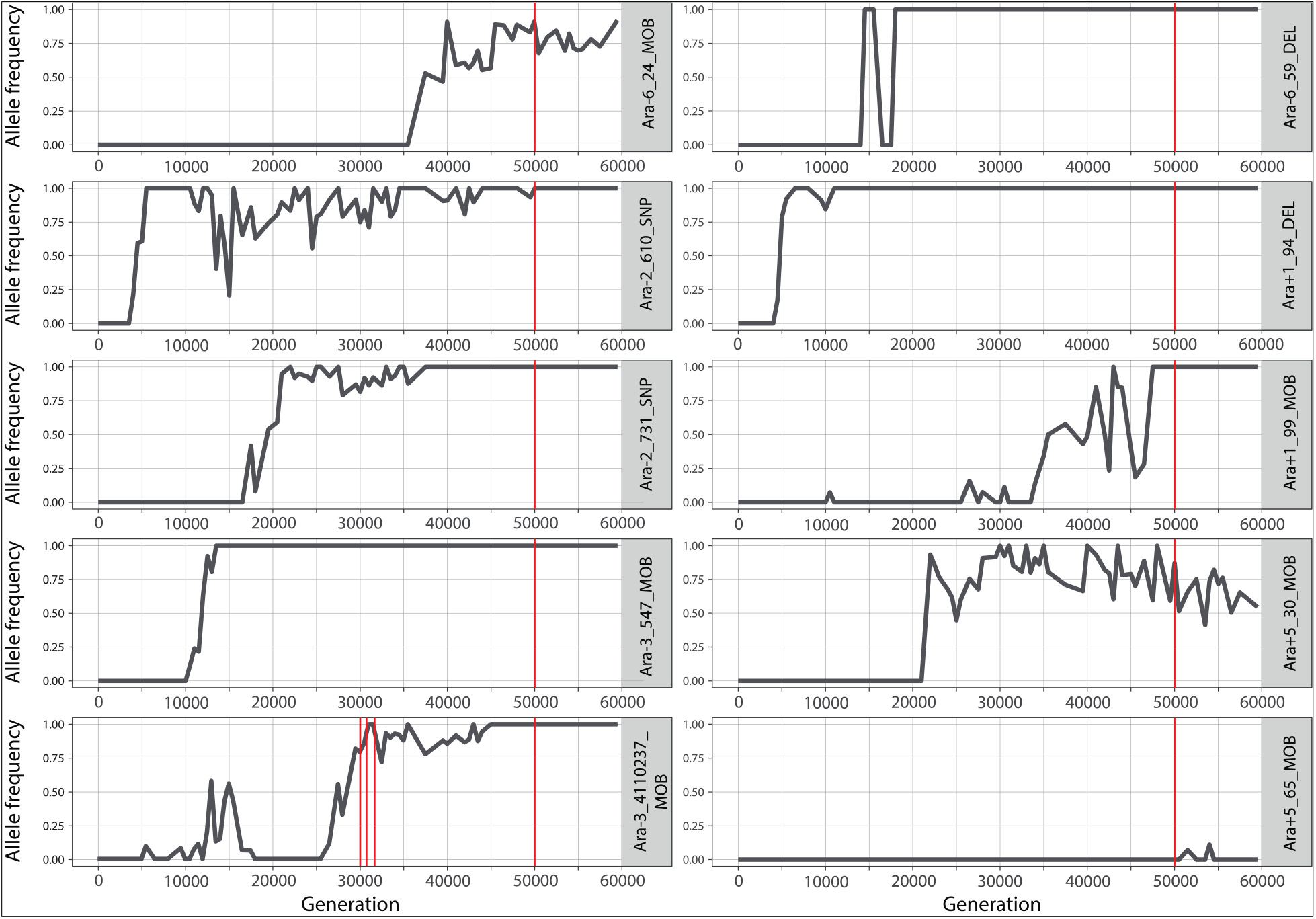
Emergence and allele frequencies of mutations associated with proto-genes. Red lines indicate timepoints at which transcription was detected.

## Discussion

Within the time-scale offered by the *E. coli* Long-Term Evolution Experiment (LTEE) [35,37], proto-genes have emerged and persisted—some in the early stages of the experiment. A primary mechanism by which proto-genes are created is through the acquisition of regulatory sequences that lead to transcription and subsequent translation of previously silent regions (Figure 1B). We observed three routes by which this occurred: (*i*) expression induced by the insertion of an IS element; (*ii*) genome rearrangements that place an existing promoter upstream of a previously non-transcribed region; and (*iii*) new mutations that create a functional promoter.

Transcription from an outward-facing promoter located at the 3’-end of IS*150* [45], due to new insertions of this element, caused nearly half of the proto-gene emergence events. IS-elements have been implicated in regulatory evolution [46,47], and in the LTEE IS-element insertions and deletions account for as many as 50% of total mutations in one population [48, 49]. IS*150* is the most actively proliferating IS-element in the LTEE; however, the increases in expression introduced by new IS150 insertions are generally modest, in most cases failing to reach statistical significance.

The most pronounced case of novel transcription was induced by the translocation of a previously silent region to a position under control of a strong promoter. This mutation, which became fixed soon after its appearance, involved a deletion in the glutamate synthase operon that placed its upstream regulatory sequences in proximity to a non-genic region. The endpoints of this deletion show no appreciable sequence similarity to one another, indicating that it was the product of illegitimate recombination. This type of event is unusual in the LTEE, since most deletions in the LTEE arise from homologous recombination between repeat elements, including IS elements [50,51].

Translocations have previously been implicated in *de novo* gene birth and the origin of new functions, for example, by generating a new sRNA gene in *Salmonella* [52] and by providing an upstream promoter to the nascent AFGP antifreeze gene in codfish [53]. Although genomic rearrangements that lead to the formation of new genes are infrequent events, relocation of the regulatory region of expressed genes can immediately confer stable transcription and translation, as required for protein-coding gene birth. As such, the large deletion in the glutamate synthase operon is the only case we observed in which the transition to a proto-gene was accompanied by translation.

Aside from promoter-capture events, proto-genes were also generated by small deletions or point mutations, although in most cases, new promoters were not evident or recognizable. SNPs and small indels represent over 95% of mutations that we observed in the LTEE, so their relatively minor contribution to the emergence of proto-genes seems puzzling given that functional promoters are pervasive in sequence space [54,55] and occur frequently in bacterial genomes [56]. However, bacteria have several mechanisms, such as H-NS [57], transcription termination factors [58], and genetic context [59], that prevent profligate and nonproductive transcription, whereas the capture of existing promoters can bypass these requirements.

Given that there is widespread and arbitrary transcription throughout the genome [14,26], initial expression of non-genic regions may be common. Whereas LTEE- evolved cells have previously been reported to contain higher mRNA abundances compared to ancestors [36], our findings also indicate that larger parts of the genome in evolved populations experience transcription. Despite the quantity of raw material for *de novo* gene formation, few non-genic regions in the LTEE are transcribed at the levels of annotated genes. For nascent transcripts to be retained by selection, expression needs to persist and reach a certain minimum threshold [38]. We demonstrate that non-genic expression persists when caused by regulatory mutations, and cannot be ascribed to stochastic transcription, as might be observed at single timepoints [15,60,61].

Based on the total number of lines examined, we estimate the rate of proto-gene emergence to be about once per 60,000 generations, with some appearing as early as 4,000 generations. To ensure that our final set of proto-genes were authentic and evolutionarily novel, we excluded all cases that did not have a readily identifiable causal mutation as well as those whose corresponding regions were expressed under non-LTEE conditions. Given the stringency of these criteria, our reported rate of proto-gene emergence can be considered a minimal estimate.

To date, most strain- and lineage-specific genes have been identified by comparative genomics, such that new ORFs represent either a transition from non-genic regions in closely related organisms or sequences gained through transfer from distantly related or unidentified taxa. In contrast, the LTEE, with the copious sequencing and transcriptomic data [35,36], permits the direct observation of proto-gene formation in individual lineages and is impervious to the horizontal transfer of genes from external sources. Genome-wide transcriptome surveys address one end of the *de novo* gene emergence puzzle, *i.e.*, the availability of raw material for selection to act upon [12]. At the other extreme, functional screens of random peptides are able to assess whether stably translated peptides confer an adaptive benefit [62–64]. The present study adopted an intermediate approach that traced how the proto-genic raw material arises, and once available, whether it persists. By combining high-resolution explorations of the genome and transcriptome, this approach has demonstrated that proto-genes continuously emerge and can contribute to the lineage-specific genes present in bacterial genomes.

## Materials and Methods

### LTEE strains

Strains were selected from the long-term evolution experiment (LTEE), which consists of 12 replicate lines of *E. coli* that have been grown in continuous culture since 1988 [65]. For the present study, we examined datasets generated for clonal isolates in: (*i*) 11 of the 12 LTEE populations (all but Ara+6) at 50,000 generations [36] and (*ii*) the Ara–3 populations at generations 5,000, 10,000, 15,000, 20,000, 25,000, 27,000, 30,000, 31,500 and 33,000. (Figure S1, Table S1). Most clones were selected because they possess sets of mutations that place them close to the lineage that evolved citrate utilization [66] (Figure S1). All datasets include comparable information for their LTEE- ancestor (REL606 or REL607).

### Transcriptomes

RNA-seq and Ribo-seq reads for the 50,000-generation clones were obtained from [36], and non-LTEE RNA-seq datasets for were acquired from [39,40]. For the Ara–3 time-series, RNA was isolated from at least three biological replicates of each strain cultured on separate days. For each replicate, we revived frozen stocks by inoculation into 10 ml of Davis Minimal media supplemented with 2 μg/l thiamine and 500 mg/l glucose (DM500). After overnight growth at 37°C, 500 μl of each culture was diluted into 50 ml of prewarmed DM500 and grown for an additional 24 hr. Subsequently, 500 μl of these preconditioned cultures were inoculated into 50 ml DM500 and grown to 30–50% of the maximum observed OD600 at stationary phase. Cells were harvested by centrifugation, washed twice with saline, and flash-frozen on liquid nitrogen.

RNA was extracted from frozen cellular pellets using RNASnap [67]. Resulting supernatants were purified using Zymo Clean & Concentrator-25 columns (Zymo Research) incorporating the on-column DNase treatment step. The integrity of purified RNA was assessed with TapeStation (Agilent), and ribosomal RNAs were depleted using the Gram-negative bacteria RiboZero rRNA Removal Kit (Epicentre). Final eluates were used as input for strand-specific RNA-seq library construction using the NEBNext RNA Library Prep Kit (New England Biolabs). Libraries were fractionated on 4% agarose E-gels (Invitrogen), and amplicons ranging from 0.2–8 kb were extracted and purified using the Zymoclean Gel DNA Recovery Kit (Zymo Research), quantified using a Qubit 2.0 fluorometer (Life Technologies), and stored at −80°C prior to sequencing. Libraries were sequenced on an Illumina HiSeq 4000 by the Genomic Sequencing and Analysis Facility (GSAF) at the University of Texas at Austin to generate 2×150-base paired-end reads. Raw FASTQ files of reads are available in the NCBI Sequence Read Archive (PRJNA896785).

### Data processing and analysis

For the datasets acquired from [39] and [40], raw reads were processed with Trimmomatic [68] by removing adapter sequences and low-quality bases from both ends, and only reads longer than 29 bases were retained. Reads from the 50,000- generation dataset were stripped of adaptor sequences, demultiplexed, dereplicated, end-trimmed based on read quality and depleted of rRNAs using scripts available from [36]. The processed FASTQ files from all datasets were mapped to their respective genomes using Bowtie2 [69], using the “local” alignment option in “very sensitive” mode, with default values for all other parameters.

### Expression analysis

Assessing gene expression changes in the context of known genomic features requires mapping reads to a reference transcriptome, a process complicated in LTEE strains because mutations have changed locations and presence in the genome. For this and all subsequent analyses, we used lists of the mutations present in each LTEE clone that were compiled in prior whole-genome resequencing studies [38,50] and are available as GenomeDiff files in the LTEE-Ecoli genomic data repository (v2.0.1) [70]. Genome sequences of each evolved clone were generated using these GenomeDiff files and the *gdtools* APPLY command in *breseq* [71].

To estimate expression in both annotated and non-coding regions of the genome, we adopted a modified version of the method used by [14]. The ancestral genome was partitioned into 400-bp windows, which were searched against each evolved genome using GMAP to extract map coordinates [72]. To account for deletions, duplications, and spurious mapping, windows that either mapped more than once, had a >10 bp insertion or deletion, overlapped with an insertion sequence, or failed to map in any of the evolved genomes in either dataset were removed from the analysis. The final list of windows common to all time-points in both datasets (*n* = 18801) covered 81.2% of the genome on both strands. Windows that overlapped with annotated genes by more than 15 bp on the same strand were marked as “annotated”, and the remainder marked as “non-genic”. This latter category was further subdivided into “antisense” and “intergenic” windows, depending on whether they overlapped a coding gene on the opposite strand.

Non-genic windows within 100 bp of an annotated gene on the same strand were considered cases of potential transcription readthrough. Overlaps and distances between sequences were determined using the “intersect” and “closest” utilities in BEDTools [73]. The transcriptome-construction and window-categorization processes are summarized in Fig S4.

For read counting and differential expression analysis, we generated separate annotation files for each LTEE clone by extracting the genome-specific coordinates of each window as informed by searches against evolved genomes using GMAP. Numbers of RNA-seq and Ribo-seq reads mapping to each feature in the annotation files were counted using the “htseq-count” tool from the HTSeq package [74]. In cases where a read mapped to more than one feature, the “nonunique-all” option of htseq-count assigned the read to both.

Normalized read-counts expressed as mean transcrsipts per million (TPM) were calculated for each replicate and averaged as follows:

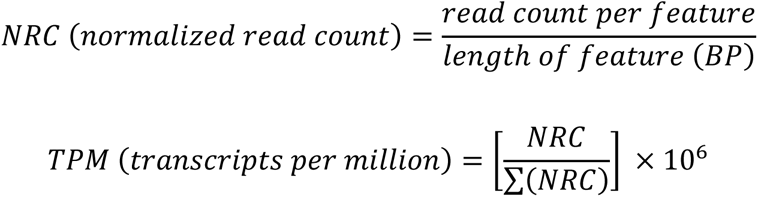

Differential expression of corresponding windows in the ancestor and each evolved line was analyzed using the DESeq2 package in R [75] with apeglm normalization [76]. A Wald test-generated *p*-value of 0.05 (adjusted by the Benjamini-Hochberg method) was used as the threshold for considering an element to be differentially expressed. To assess differential translation of windows while accounting for changes in transcription, we used the Riborex package [77] according to scripts provided in [36], with a *q*-value of 0.01 used as the threshold for significance.

### Proto-gene detection and characterization

To identify increases in transcription associated with the appearance of new mutations in the 50,000-generation dataset, we extracted 100- and 200-bp regions immediately downstream of mutations from each ancestral and evolved genome, under the expectation that changes in expression would occur within this distance. For mutations other than those caused by IS*150* insertions, which have promoters oriented on a particular strand, regions were extracted from both strands. We then removed regions with >10-bp same-strand overlap with any annotated RNA gene, protein-coding gene, pseudogene, or repeat region with the “intersect” utility of BEDTools. To generate normalized read counts for later steps, we added in annotated gene coordinates to this list of sequences to construct our final transcriptome to identify proto-gene emergence. Read counting and differential expression analysis were conducted as described above.

All regions exhibiting statistically significant increases in transcription relative to the ancestor were extracted and mapped back to their respective genomes with GMAP. Regions appearing more than once, cases where the adjacent mutations are counted twice by the breseq pipeline, and those that overlap with repeat regions on the opposite strand were removed, leaving a total of 63 regions with downstream mutations with increased expression in the 50,000-generation dataset. For these 63 regions, we generated transcription coverage plots, which were visually inspected for the presence of ancestral transcription. To accomplish this, we converted each bam file into a genome coverage file with the “genomecov” utility in BEDTools, extracted coverage information for each region of interest, and visualized them with the ggplot2 package in R [78]. Of the 63 initial cases, 38 were excluded as containing ancestral transcription, with a further 5 being classified as “ambiguous” (Table S4). Proto-genes were extracted from the time-series data in an identical manner, with minor modifications on account of the paired-end dataset available for this series.

To determine if candidate regions exhibit transcription under conditions that differ from those in the LTEE, we leveraged two large RNA-seq datasets generated from the ancestral LTEE strain grown in a variety of environments [39,40]. After processing raw files, we produced read counts and coverage plots for the 19 proto-gene candidates in each of the 152 RNA-seq samples, as described above (Figure S3). We also searched for occurrences of the candidate proto-genes in the K-12 MG1655 transcriptome reported in [41], which was assembled from 3,376 RNA-seq datasets deposited in the Sequence Read Archive [79]. We extracted the co-ordinates of all annotated and non-annotated transcripts from the associated supplementary tables, extracted their sequences, and used blastn [80] to query this database with proto-gene sequences.

All proto-genes passing these filters were checked for the presence of newly formed promoters with iPromoter-2L [43] and the Promoter calculator [44]. As evidence of potential translation, we searched for Ribo-seq reads within transcribed regions and constructed coverage plots as described above. To determine the occurrence and frequency of proto-gene-causing mutations in the LTEE, we used the mixed-population sequencing data generated from each preserved strain in the LTEE (separated by 500- generation intervals) from the first 60,000 generations [35]. After trimming the raw files with fastp [81], we searched each time point with breseq to identify mutations responsible for the proto-genes. We extracted mutation frequencies in each population from the breseq-generated GenomeDiff files, and visualized them with the ggplot2 package in R. Scripts used for analyses conducted in this study are available at https://github.com/Hassan-1991/LTEE-new-gene.

## Acknowledgements

We thank Kim Hammond for figure preparation, and Drs. Zachary Ardern and Daniel Deatherage for their helpful comments. This work was supported by the U.S. National Science Foundation (DEB-1951307 to J.E.B), the U.S. Army Research Office (W911NF- 12-1-0390 to J.E.B.) and the National Institute of Health (R35GM118038 to H.O.). The funders had no role in study design, data collection and analysis, decision to publish, or preparation of the manuscript. J.E.B. and H.O. received summer salary from the U.S. Army Research Office and the National Institute of Health grants, respectively; and M.U. received academic-year stipend from the National Institute of Health grant.

## Supporting information

**Supplementary File 1. Windows displaying transcription, translation, and differential expression in 50,000-generation populations in the LTEE.**

**Supplementary File 2. Coverage plots of ancestral and evolved transcription in regions with upstream mutations that showed significantly higher expression in the 50,000-generation.** Purple and orange lines represent ancestral and evolved transcription, respectively.

**Supplementary File 3. Coverage plots of transcription in regions with upstream mutations that showed significantly higher expression in any of the evolved time points in the time-series dataset.**

**Table S1. Strains used in this study.**

**Table S2. Mutation-linked novel transcription identified in both datasets. Table S3. Mutations in non-genic regions investigated in the study.**

**Fig S1. Relationships among clones in the time-series dataset.** All but two clones cannot utilize citrate: ZDB564 has a rudimentary (Cit+) and CZB154 has a fully developed (Cit++) phenotype. Two clones (ZDB199, ZDB200) stem from highly diverged clades that did not evolve citrate utilization. Figure adapted from [66].

**Fig S2. Transcription of non-genic regions at different time-points in the Ara–3 population.** Comparisons of ancestral and evolved populations showing numbers of non-genic windows with normalized RNA-seq read counts expressed in average number of transcripts per million (TPM).

**Fig S3. Coverage plots depicting transcription of candidate proto-genes in the non-LTEE expression library.** The *y*-axis ranges are set according to the expression level of proto-gene candidates in the LTEE. Only cases having between 3 and 50 reads in at least one condition are shown.

**Fig S4. Transcriptome construction and categorization strategy for 400-bp windows in the 50,000-generation dataset.**

